# Infra-slow (<0.1 Hz) modulation of human brain pulsations in awake and sleep states

**DOI:** 10.64898/2025.12.15.692942

**Authors:** Tommi Väyrynen, Heta Helakari, Vesa Korhonen, Johanna Tuunanen, Niko Huotari, Janne Kananen, Seyed-Mohsen Ebrahimi, Ahmed Elabasy, Matti Järvelä, Katariina Hautamäki, Katariina Laurén, Lauri Raitamaa, Ulla Salmi, Johanna Piispala, Mika Kallio, Vesa Kiviniemi

## Abstract

Human brain exhibits three propagating pulsations, namely cardiovascular, respiratory, and vasomotor waves, which together propel the flow of intracranial fluids. While their pulsation characteristics have been extensively studied, their causal interconnections have not been systematically investigated. Using ultrafast whole brain magnetic resonance encephalography (MREG), we analysed the frequency domain up to 5 Hz for cross-frequency oscillatory interactions in awake and NREM-sleep states of 23 healthy volunteers. Using transfer entropy (TE) analysis, we found that in the awake state the infra-slow (ISF < 0.1 Hz) oscillations of statistically independent resting state networks (RSN) largely drove the neurofluid (NF) pulsatility. NREM-sleep was associated with increased power of infra-slow fluctuation (ISF) vasomotor oscillations and with altered driving patterns between RSN and NF networks in the direction of a causally chained pattern. Importantly, within these independent signal sources, we found three distinct cross-frequency coupling frequency ranges occurring at ISF (<0.1 Hz), respiratory (∼0.25 Hz), and cardiovascular (∼1 Hz) frequencies, where the slower pulsations generally modulated the faster ones, except for a finding of inverted cardiorespiratory drive in NREM-2 sleep. These results indicate the presence of directional ISF-coordinated mechanisms underlying brain pulsations that contribute to driving the intracranial fluid transfer processes.

**Significance statement:** Cerebrospinal fluid (CSF) flow is essential for brain fluid homeostasis and interstitial metabolite clearance. Human brain exhibits three types of intracranial pulsations linked to CSF flow, which are particularly distinct during sleep, when fluid clearance processes are most active. We predicted that these pulsations, despite their independent sources, interact with each other to coordinate CSF flow. Using functional magnetic resonance imaging (fMRI) during wakefulness and non-rapid eye movement sleep (NREM), we investigated cross-frequency coupling patterns up to 5 Hz within the brain. Results revealed a novel mechanism in human brain whereby infra-slow (ISF) vasomotor oscillations coordinated faster brain pulsation dynamics, which could be a factor mediating the increased perivascular clearance during sleep.

## Introduction

Human brain relies on circulation of cerebrospinal fluid (CSF) through low resistance perivascular spaces, allowing the influx of CSF into brain cortex and mixing with the interstitial fluid (Iliff et al., 2012). This fluid transport is vital for removal of metabolic waste products, which is a prerequisite for normal brain function (Kelley and Thomas, 2023). Sleep associated increase of the interstitial space between neurons and astrocytes, promotes more efficient CSF flow (Xie et al., 2013). Conversely, decreased sleep duration has been linked to accumulation of amyloid-β (Shokri-Kojori et al., 2018).

Considerable effort has been put into understanding the physiological drivers of net CSF flow in brain parenchyma. Three types of intracranial pressure fluctuations: infra-slow (ISF) vasomotor (∼0.02 Hz), respiratory (∼0.25 Hz), and cardiovascular pulsations (∼1 Hz), oscillate within human brain, and can be detected with rapidly sampled fMRI magnetic resonance encephalography (MREG) sequence. All three pulsations are thought to independently contribute to net CSF flow by mechanically displacing perivascular CSF (Hadaczek et al., 2006; Iliff et al., 2013; Dreha-Kulaczewski et al., 2015, 2017; Mestre et al., 2018; van Veluw et al., 2020; Bojarskaite et al., 2023; Hauglund et al., 2025).

Initial studies suggested that cardiovascular pulsatility drives the inflow of subarachnoid CSF to the cerebral cortex of mice (Hadaczek et al., 2006; Iliff et al., 2013; Mestre et al., 2018). Other research lines suggested that downward flow of venous blood during inspiratory phase of respiration is counterbalanced by upward flow of CSF towards head (Dreha-Kulaczewski et al., 2015, 2017; Kiviniemi et al., 2016).

Most recently, arterial vessel wall oscillations, known as vasomotor waves, have been proposed to propel brain CSF flow (van Veluw et al., 2020; Bojarskaite et al., 2023; Hauglund et al., 2025). Optogenetic stimulation of noradrenergic neurons of locus coeruleus evoked anticorrelated vasomotion and CSF flow in mice (Hauglund et al., 2025). Similarly, optogenetic stimulation of arterial oscillations enhanced CSF inflow, indicating that vasomotion contributes to pumping CSF into the brain. By current understanding, the controlled release of the vasoconstrictor norepinephrine (NE) from ascending projections of the locus coeruleus coordinates widespread vasomotor waves during sleep (Osorio-Forero et al., 2021; Kjaerby et al., 2022; Lüthi and Nedergaard, 2025).

In humans, vasomotor-induced blood volume fluctuations are also accompanied by opposing changes in cortical CSF volume during non-rapid eye movement (NREM) sleep stages 1 and 2 (Borchardt et al., 2021; Väyrynen et al., 2024). Moreover, the oscillation power and propagation speed of these waves both increase during NREM sleep (Helakari et al., 2022; Elabasy et al., 2025), and further synchronize the arterial vasomotor activity with concomitant venous BOLD signal (Tuunanen et al., 2024). Disruption of the characteristics of brain pulsations occur in several neurological conditions. For example, patients with type 1 narcolepsy show increased vasomotor waves, but reduced cardiovascular pulsatility in awake state compared to healthy controls (Järvelä et al., 2025). Diverse other central nervous system disorders including Alzheimer’s disease, epilepsy, and lymphoma have been linked to changes pulsation dynamics (Kananen et al., 2020, 2022; Tuovinen et al., 2020; Rajna et al., 2021; Poltojainen et al., 2022).

These clinical associations and mice experiments draw attention to the possibility that the three physiological pulsations may be synchronized. The limited temporal resolution of conventional echo planar imaging BOLD sequences constrains the detectable frequency range, making it difficult to capture fast and slow oscillatory components simultaneously (Huotari et al., 2019). As a result, there has been no systematic investigation of the hemodynamic coupling patterns in human brain.

To address this issue, we used a T2*-based fMRI MREG sequence sampled at 10 Hz to accurately capture pulsation dynamics. We searched for cross-frequency coupling patterns using phase-amplitude coupling estimator (PAC) (Canolty and Knight, 2010) during EEG-verified awake and NREM1-2 sleep in healthy volunteers. We further employed phase transfer entropy (TE) to estimate the information transfer between the coupled frequency ranges (Lobier et al., 2014), testing our hypothesis that ISF vasomotion would coordinate faster cardiorespiratory oscillations.

## Results

### Infra-slow phase is coupled with faster amplitude oscillations in human brain

To establish the frequency specific coupling among pulsations in human brain, we quantified PAC in the MREG signal across the frequency domain extending from 0.01 Hz to 5 Hz. Voxel level analysis revealed three separate clusters deviating from baseline PAC (Fig. 1c) and involving ISF, respiratory, and cardiac frequencies, which reached maximal coupling during NREM-2 sleep. The first coupled frequency range involved ISF amplitudes, which were coupled with even slower ISF phase changes: f_amp_(0.02-0.1Hz); f_phase_(0.01-0.02 Hz). The second coupling range matched respiratory amplitudes, which were similarly coupled with ISF phase changes: f_amp_(∼0.25 Hz); f_phase_(0.01-0.05 Hz). The third coupling frequency range matched cardiac amplitudes, which were coupled with a wider phase bandwidth: f_amp_(∼1 Hz); f_phase_(0.01-0.1 Hz).

**Figure 1.**
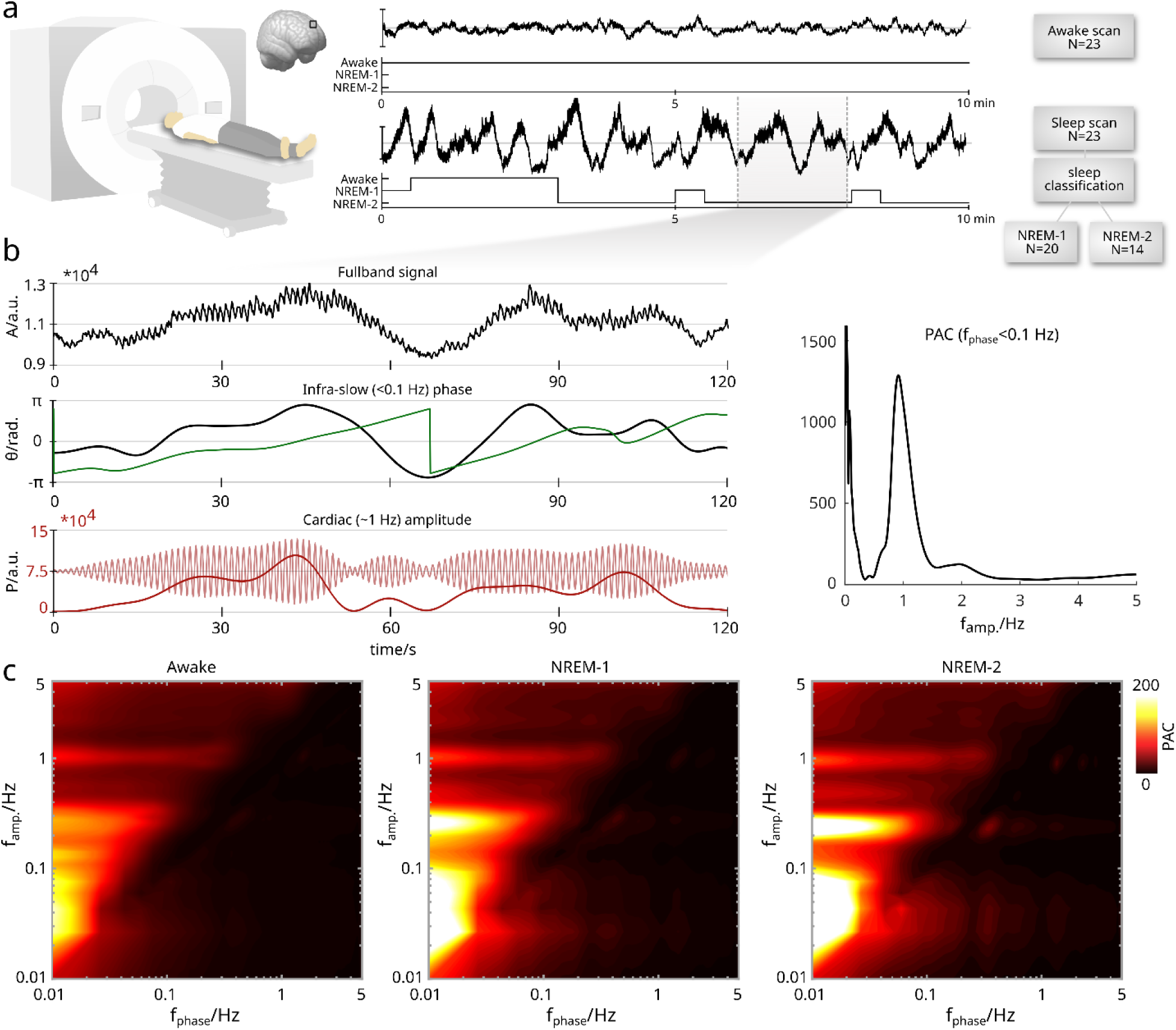
Infra-slow fluctuation (ISF, <0.1 Hz) signal changes are coupled with physiological pulsation powers. a) Example signal for an entire 10-minute sequence scan time during awake and sleep sessions, along with EEG-derived hypnograms. b) Top row illustrates a two-minute signal epoch during NREM-2 sleep, where phase-amplitude coupling (PAC) of ISF phase and cardiac amplitudes (∼1 Hz) occurred. The middle row shows the ISF bandpass filtered signal and the corresponding phase time-series. The bottom row shows the cardiac frequency filtered signal and associated cardiac power fluctuations over time. The right middle row shows PAC of the ISF phase to amplitudes ranging from 0.01 to 5 Hz, which confirms the visual observation that ISF phase couples with cardiac power around 1 Hz. c) Average whole brain PAC estimates across frequency space in awake, and NREM-1 and NREM-2 states, reveal three coupling frequency ranges, which increase in magnitude as a function of sleep stage. The x-axis illustrates the phase frequency, whereas the y-axis indicates the frequency of amplitude, and color scale represents the PAC coupling magnitude.

The coupling especially of respiratory and ISF amplitudes demonstrated higher mean PAC levels in comparison to cardiac amplitudes. NREM-1 and -2 sleep stages were both associated with elevated average PAC values in contrast to awake state. Confirming our hypothesis, we found multiple coupled frequency bands in the hemodynamic signal, which corresponded to the well-known brain pulsation frequency ranges: ISF, respiratory, and cardiac range.

### Sleep increases infra-slow fluctuations of RSN and NF components

It has been unknown how the three brain pulsations are expressed in independent functional units of the human brain. To quantify power contributions of simultaneous cardiorespiratory and vasomotor effects on neurofluid (NF) and on functionally connected brain networks, we computed the spatially independent components (IC) from the MREG data. The ICs were categorized into NF components and classical resting state networks (RSN), after visual inspection by an expert neuroradiologist (V.Ki.) (Fig. 2 a,d, Supplementary Table 1). We then assessed the components frequency domain by analyzing signal power levels up to 5 Hz in awake and sleep states.

**Figure 2.**
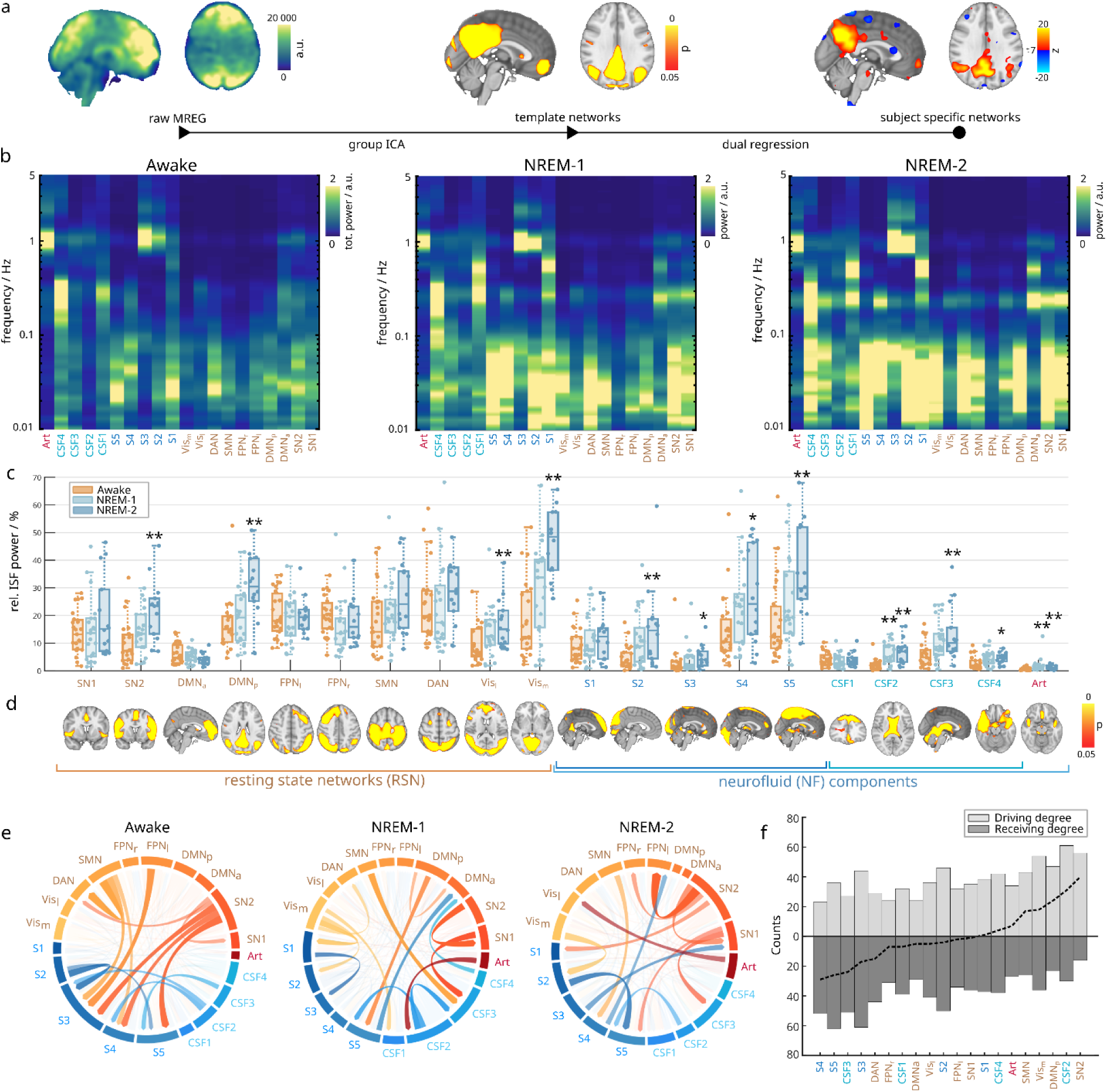
Spectral characteristics of resting state networks (RSN) and neurofluid (NF) components of brain. a) Group independent component analysis (ICA) was used to separate RSN from spatially independent NF sources. b) Average ICA spectra are depicted for the 0.01-5 Hz frequency band in awake and NREM-1/2 sleep states. c) Infra-slow (<0.1 Hz) powers relative to total signal power levels, with units in percentiles. Awake state was compared to NREM-1 and -2 states, where statistical significance is denoted by asterisks (*p_adj_*< 0.05*, 0.01**, 0.001***). d) Spatial maps of ICA template networks in the most representative horizontal, coronal, or sagittal slice, with assignment of visually labeled components to classical RSN (left) and NF (right) components. e) Prediction patterns between the ICs, measured by phase transfer entropy (p<0.05) is visualized in network diagrams, where the strongest connections are displayed. The arrows visualize the directions of interactions. f) The degree of coupling is pictured, where numbers of significant links are displayed on the ordinate axis.

As expected, fast cardiovascular pulsations were prominent in the IC labelled as the cerebral artery Art, but were also strongly expressed in venous sinus S2 and S3 components (Fig. 2b). In contrast to NF components, RSN components in general exhibited less cardiovascular activity, except for the anterior default mode network DMN_a_ and salience network SN2 located close to the anterior and medial cerebral artery branches. Slower respiratory fluctuations were most dominant within CSF spaces CSF1,3,4, but also visible in the venous S1 and S4 components, and in the lateral visual Vis_l_ and SN2 networks. The ISF powers were highly pronounced especially in venous S1, S4, S5, and with RSNs in the dorsal attentional network DAN, DMN_p,a_ and SN2. Notably, the ISF power was distributed across a broad frequency range below 0.1 Hz. The IC components showed unique ISF characteristics, with a patchy appearance rather than having a single frequency centered band.

We then proceeded to make a statistical assessment of the ISF powers relative to total signal power levels in awake and NREM1-2 sleep states (Fig. 2c, Supplementary Table 2). Based on our previous voxel level studies (Helakari et al., 2022; Väyrynen et al., 2024), we hypothesized that ISF power would increase in the collective ICs during NREM-sleep. The overall levels of ISF band powers were higher in RSNs (P_A_=15.7%, IQR=8.8%; P_N1_=17.6%, IQR=9.9%; P_N2_=22.7%, IQR=7.2%) in comparison to NF-labeled components (P_A_=5.9%, IQR=4.9%; P_N1_=9.9%, IQR=7.2%; P_N2_=12.3%, IQR=7.6%) (Fig 2c). Eight of the ten NF components showed significantly increased ISF power during sleep. In particular, the arterial component showed increased ISF bandpower in comparison to both NREM states (Art: A-N1: *Z*=-3.7, *p_adj_*<0.01**, A-N2: *Z*=-3.2, *p_adj_*<0.01**), even though absolute levels of ISF powers remained low and dominated by cardiac activity. All CSF components except for the frontal CSF1 component had increased power during sleep (CSF2: A-N1: *Z*=-3.8, *p_adj_*<0.01**, A-N2: *Z*=-3.5, *p_adj_*<0.01**). Similarly, all venous sinus components except for frontal S1 also showed increased oscillation power in the ISF range (S5: A-N1: *Z*=-1.7, *p_adj_*=0.1, A-N2: *Z*=-3.1, *p_adj_*<0.01**).

Somewhat surprisingly, only four among the ten classical RSNs studied showed significant increases in ISF bandpower during sleep (Fig 2c). Both visual networks: lateral and medial increased ISF power (Vis_m_: A-N1: *Z*=-2.0, *p_adj_*=0.08, A-N2: *Z*=-4.2, *p_adj_*<0.01**). Also, posterior parts of DMN showed increased ISF power in NREM-2 sleep (DMN_p_: A-N1: *Z*=-0.9, *p_adj_*=0.5, A-N2: *Z*=-3.6, *p_adj_*<0.01**), but anterior parts did not. In the salience network, SN2 but not SN1 showed increased power (SN2: A-N1: *Z*=-2.1, *p_adj_*=0.08, A-N2: *Z*=-3.4, *p_adj_*<0.01**). The dorsal attentional DAN, sensorimotor SMN and frontoparietal FPN_l,r_ networks also showed no increase in ISF power levels. See Table S2 for full list of statistical results.

In line with our hypothesis, ISF power increased during sleep, encompassing NF components and especially posterior RSNs, possibly reflecting increased fluctuations in cerebral blood flow caused by increased vasomotor wave activity.

### Connections between RSN and NF components are brain state dependent

Although the functional connectivity between the RSNs have been rigorously studied over the years, it remains unknown how the NF components relate to the classical RSNs. As the ISF changes in T2* weighted BOLD fMRI signal mainly reflects relative de/oxyhemoglobin changes in post-capillary veins, it follows that flow changes in the NF, especially venous components, could potentially affect RSN signal changes. To further map the connections between NF components and functional RSNs, we calculated TE between ICs in the ISF range (Fig. 2e, Supplementary Fig. S1).

Extracting significant connections (p<0.05) between the ICs revealed that RSNs generally predict NF components (Fig. 2e). TE patterns were most coherent in the awake state, where especially the SMN and SN2 networks predicted the venous sinus components S3-5. Additionally, the SMN and SN2 networks also predicted DAN and SN1, whereas CSF components CSF2-3 predicted major venous sinus outflow components S3-5.

In NREM-1 sleep, there was no longer a stable RSN prediction of NF, and the status of SMN and SN2 predictors changed into more complex interactions. An interesting physiological chain effects could also be identified during sleep: the cerebral arterial component Art started to predict CSF in the lateral ventricles CSF2, which, together with SN2 and SMN network activity, predicted downstream 3rd ventricle and aqueduct signal in the CSF3 component. The posterior DMN_p_ and medial visual Vis_m_ networks both predicted changes in posterior sinus S2 and fronto-parietal sagittal sinuses S5, which further together with perisylvian CSF4 formed a connection back to anterior parts of the DMN_a_. Posterior DMN_p_ was also directly connected to the anterior DMN_a_, whereas the medial visual Vis_m_ network also predicted the frontal cortex CSF1 component and DAN.

In NREM-2 sleep, two frontal drivers emerged: SN2 and SN1. SN2 predicted both FPN_l,r_ and SN1 networks, and among the NF components also the 3rd ventricle and aqueductal CSF3 and the upper sagittal sinus S5 components. Much as in the awake state, SMN predicted S4 venous changes. Vis_m_ continued to predict venous sinus S2, as in lighter NREM-1 sleep. Also, the arterial component predicted the lateral Vis_l_ network.

To determine which networks were most actively contributing to overall network dynamics, we calculated the degree of coupling, which refers to the number of links to other networks (Fig. 2f). We further separated the degree of coupling degree into driving and receiving, thus considering the interaction directionality. Results revealed that SN2, ventricular CSF2, DMN_p_, Vis_m_, SMN and arterial components were most actively involved in network driving dynamics, having a net positive effect. In contrast, the venous sinuses S3-5, CSF3 and DAN were especially associated more as receiving nodes, apparently following the dynamics of other networks rather than being coordinators.

Effective connectivity results showed the ISF RSN phase transitions to be predicting NF components rather than other way around. The RSN prediction was most coherent during wakefulness, whereas sleep reorganized these functional connections. In line with this, venous sinuses were characterized as receiving nodes, reacting to changes in RSNs activity.

### Independent brain RSN and NF sources reveal three cross-frequency coupling frequency ranges

Using statistically independent signal sources, we re-analyzed the coupling patterns observed at the whole brain level. We tested the hypothesis that the RSNs and NF components would exhibit distinct coupling patterns, reflecting their different signal origins. Our analysis of ICs revealed three statistically significant (p < 0.05) cross-frequency coupling ranges (Fig. 3a). In two of these frequency ranges, the ISF phase was involved. The first such frequency range exhibited coupling between the ISF phase (∼0.02 Hz) and slow-frequency (SF) amplitudes (∼0.04–0.11 Hz), which was especially conspicuous during NREM-2 sleep (labeled as “1.” Fig 3a.). The second such range was also most prominent during NREM-2 sleep and involved a wider ISF phase band (∼0.01–0.05 Hz) coupled with respiratory amplitudes (∼0.25 Hz) (labelled as “2” Fig 3a). The third coupling range was observed across both wakefulness and sleep and involved respiratory phase (∼0.25 Hz) coupling with cardiac amplitudes (∼1 Hz). Henceforth, we refer to these cross-frequency interactions as: 1. ISF–SF, 2. ISF–Resp, and 3. Resp–Card interactions.

**Figure 3.**
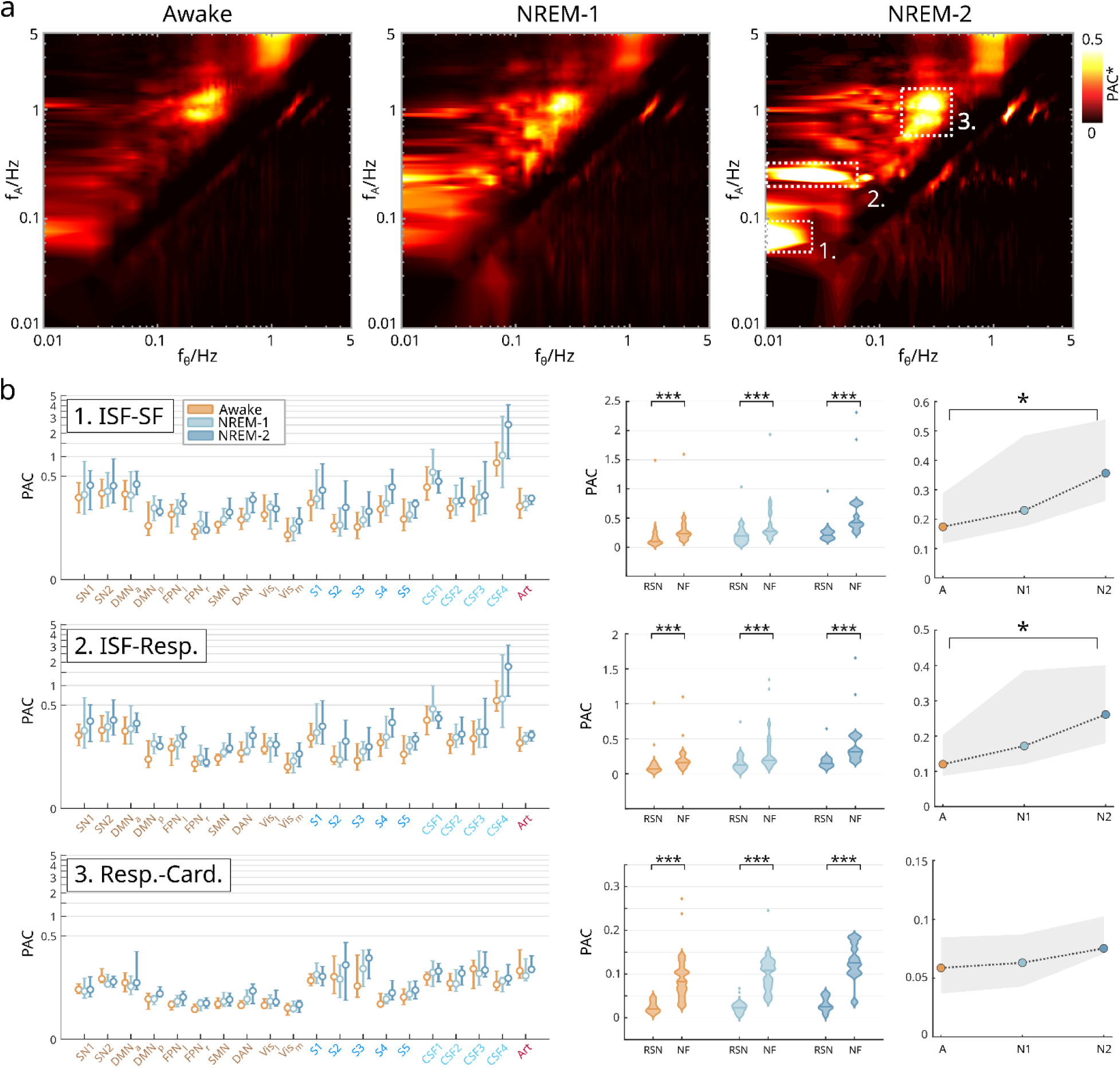
Significant cross-frequency coupling ranges associated with independent components. a) The images illustrate frequency-frequency maps of significant (p<0.05*) phase-amplitude coupling (PAC) values between phase (x-axis) and amplitude (y-axis) frequencies across wakefulness and sleep stages. b) Unthresholded PAC values extracted for each coupling ranges (as numbered 1.-3.) and for each IC. The middle-tiles show the effects of resting state networks (RSN) and neurofluid (NF) components averaged over all components. The right-hand tiles show the grand average PAC over for wakefulness and sleep states, indicating increasing coupling with sleep stage. Statistical significance is denoted by asterisks (*p_adj_*< 0.05*, 0.01**, 0.001***)

An in-depth analysis of these three coupling frequency ranges revealed similar PAC structure among components with ISF–SF and ISF–Resp couplings (Fig. 3b, left). Among the RSN components, increased PAC relative to baseline was observed in SN1, SN2, and DMN_a_, whereas among NF components, high PAC was found in venous S1, S4 and CSF-related components CSF1, CSF4. With Resp–Card coupling, PAC values were higher especially in NF components and to a lesser extent in RSNs.

With all three coupling patterns we found that the NF components had significantly higher (p<0.001***) coupling values in comparison to the RSN (Fig. 3b, middle). This effect was seen robustly across arousal states. See supplementary Table 3 for statistical analysis. Moreover, we found that sleep was also associated with increased PAC values in the ISF-SF and ISF-Resp. couplings (Fig. 3b right). With ISF-SF coupling, the median PAC was 0.174 (IQR=0.172) in awake state, 0.230 (IQR=0.309) in NREM-1 sleep, and 0.356 (IQR=0.277) in NREM-2 sleep. The difference in medians between awake and NREM-2 sleep was statistically significant (*Z*=-2.83, *p_adj_*<0.05*). Similarly, ISF-Resp. coupling during awake state had a median PAC 0.120 (IQR=0.117), which increased to 0.172 (IQR= 0.265) in NREM-1 sleep, and 0.261 (IQR= 0.221) in NREM-2 sleep, where the difference between awake-NREM-2 was statistically significant (*Z*=-2.83, *p_adj_*<0.05*). Full results are presented in supplementary Table 4.

As expected, we found three cross-frequency coupling ranges at the component level, which deviated from voxel level analysis. Surrogate statistics ensured unbiased PAC estimation, which otherwise could be attributed to underlying signal properties or to random effects. Sleep state was associated with increased coupling values.

### Infra-slow network dynamics coordinate slow oscillations and respiration

After identifying the frequency bands associated with the coupling patterns, a key question remained: in which direction do these interactions take place? To answer this, we further investigated the newly-identified coupling frequency ranges by assessing the transfer entropy (TE) within the interactions (Figure 4a). We hypothesized that the observed PAC coupling patterns would be coordinated by the slower oscillations.

**Figure 4.**
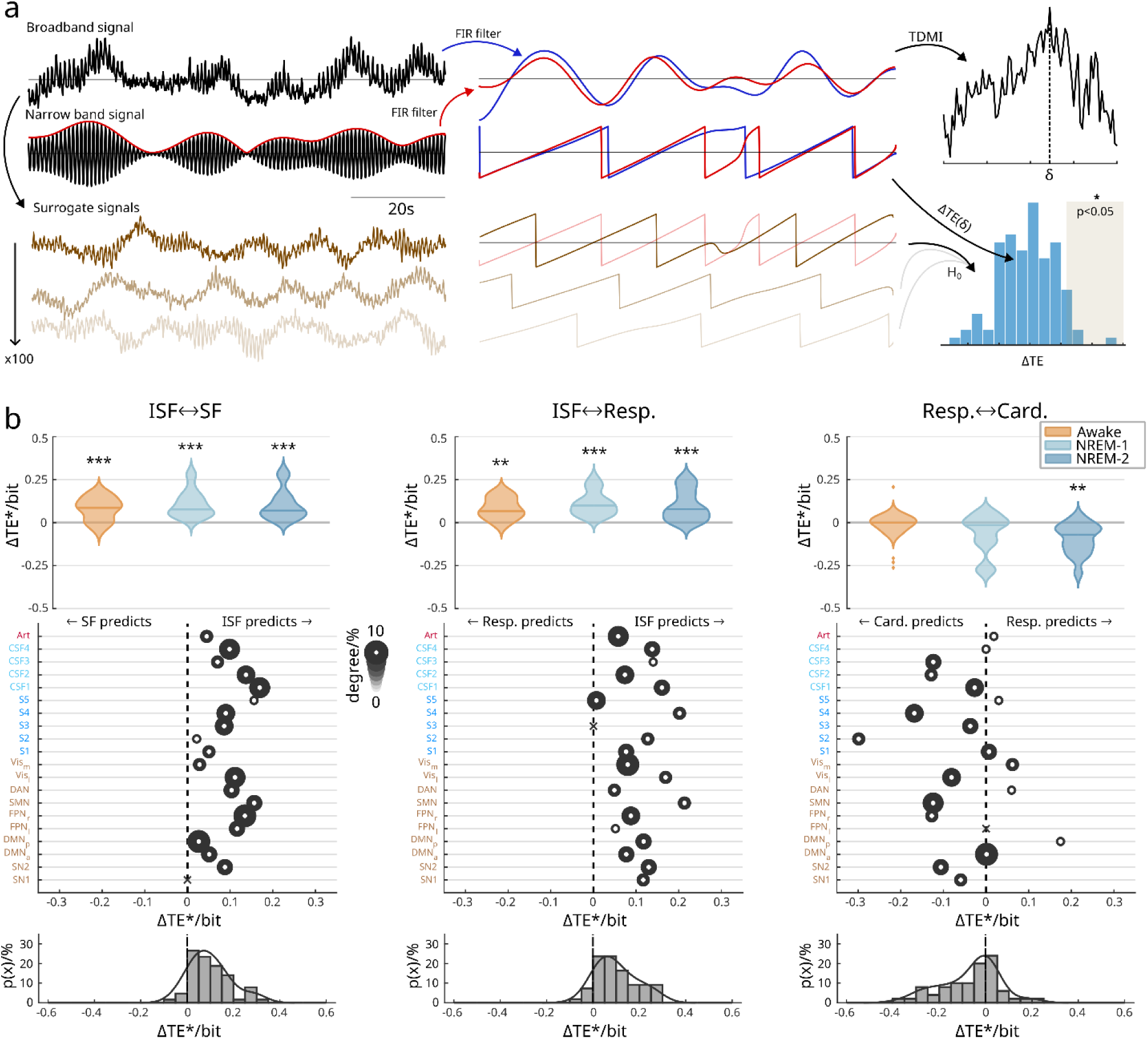
Infra-slow (ISF) network dynamics predict slow and respiratory frequency oscillations. a) In the procedure for directional inferences, we first filtered the broadband signal into two target narrowband frequencies. The phase was extracted from the lower-frequency component, while the analytic envelope of the higher-frequency oscillations was computed and subsequently filtered at the corresponding lower frequency. Phase time series were then derived, and the optimal analysis lag was determined using time-delayed mutual information (TDMI). This lag was subsequently applied in estimating directional interactions via phase transfer entropy (TE). Statistical significance was assessed using surrogate data distribution generated from the broadband signal by the time-shift method. b) Violin plot illustrates significant (p<0.05) *ΔTE* between 1. ISF-SF, 2. ISF-Resp and 3. Resp-Card coupling patterns in awake, and NREM-1 and NREM-2 sleep states. Below, the scatter plot depicts the contribution of each independent component (IC) as an average over subjects. The size of the circle represents the number of significant values in percentiles. At the bottom, probability density estimate collapsed over the IC, demonstrating that on average the ISF coordinates SF and Resp. fluctuations, while in Resp-Card coupling, cardiac pulsation is actually the net predictor of respiratory oscillations. Statistical significance is denoted by asterisks (*p_adj_*< 0.05*, 0.01**, 0.001***).

To test this hypothesis, we first evaluated the null hypothesis that the distributions of *ΔTE* would be centered around a zero median, which would suggest no net directionality. Contrary to this hypothesis, we found significant directional information transfer in the ISF-SF (*ΔTE_A_=0.084 bit, p_adj_<0.001***; ΔTE_N1_=0.078 bit, p_adj_<0.001***; ΔTE_N2_=0.070 bit, p_adj_<0.001\*\*\**) and ISF-Resp (*ΔTE_A_=0.067 bit, p_adj_<0.01**; ΔTE_N1_=0.101 bit, p_adj_<0.001***; ΔTE_N2_=0.076 bit, p_adj_<0.001\*\*\**) couplings across all arousal states (Fig. 4b). In both the ISF-SF and ISF-Resp coupling frequency ranges, the ISF phase acted as the net predictor of faster frequencies. In contrast, for the Resp-Card coupling, a significant net directionality was observed only during NREM-2 sleep (*ΔTE_A_=-0.0004 bit, p_adj_=1.0; ΔTE_N1_=-0.012 bit, p_adj_=1.0; ΔTE_N2_=-0.072 bit, p_adj_<0.01\*\**) (see Supplementary Table 5).

Arousal state had little effect on the *ΔTE* magnitude; the ISF–SF and ISF–Resp couplings did not show notable changes across wakefulness and sleep stages. However, the Resp–Card interaction showed no net directionality during wakefulness and NREM-1 sleep, but shifted to negative *ΔTE* during NREM-2 sleep, indicating cardiac prediction over the respiratory activity.

All ICs demonstrated consistent prediction directions in the ISF–SF and ISF–Resp couplings. Only small differences were observed between RSN and NF components, suggesting equal expression of coupling in the studied ICs. In contrast, the Resp–Card coupling displayed larger differences between components. While most components displayed cardiac amplitude prediction of respiratory phase, DMN_p_, Vis_m_, and DAN showed respiratory prediction over cardiac changes.

Overall, our results generally support a slow to fast hierarchical organization of human brain pulsations, as evidenced by the observed directional net prediction patterns. However, contrary to our expectations, cardiac amplitude fluctuations during NREM-2 sleep predicted respiratory changes.

## Discussion

In this study we investigated for the first time the coupling of the three major physiological brain pulsations—vasomotor, respiratory, and cardiovascular waves. Using 10 Hz MREG fMRI, we discovered that these brain wide pulsations are not independent, but are rather co-modulated through cross-frequency coupling. Specifically, we identified three frequency specific coupling frequency bands: ISF to slow (ISF-SF), ISF to respiratory (ISF-Resp), and cardiac to respiratory (Card-Resp) couplings. Along with increasing power also the coupling magnitudes of the pulsations increased across wakefulness to NREM-1 and -2 sleep states. Functionally the cortical RSN networks predominantly drive NF compartments in awake state but in sleep the ISF power increases and the causal connections chain up and form more bi-directional cascade patterns.

### Hierarchical organization of human brain pulsations

Our results showed that ISF vasomotor waves modulated the amplitude of slow oscillations (ISF-SF) and that the broader ISF band also modulated respiratory oscillations (ISF-Resp). The respiratory phase fluctuations were in turn coupled with (Resp-Card) cardiac amplitude. Phase transfer entropy analysis further revealed information transfer from slow to faster rhythms, confirming that the ISF phase acted as a coordinator of higher-frequency pulsations. Similar hierarchical slow-to-fast organization occurs with many physiological systems for example circadian rhythms coordinating hormone release (Gnocchi and Bruscalupi, 2017), the respiratory cycle influencing heart rate in respiratory sinus arrhythmia (Yasuma and Hayano, 2004), and slow electric oscillations in brain modulating fast cortical processing (Väyrynen et al., 2023).

Although our results largely support a slow-to-fast model of hierarchical organization of brain pulsations, cardiorespiratory coupling deviated from this general rule. Others have established that cardiorespiratory coupling can have bidirectional dynamics, with various physiological regulation mechanisms within the thorax (Barnett et al., 2021). Here, awake and NREM-1 states TE demonstrated no net interaction direction in either way, but cardiac pulsatility during NREM-2 sleep started predicting respiratory fluctuations. This suggests the occurrence of altered cardiorespiratory coupling during deep sleep, with reversed fast-to-slow modulation of cardiorespiratory coupling.

### Component-level insights into neurovascular coupling

The BOLD contrast in the low-frequency range reflects a mixture of signal sources that vary by spatial location. In this frequency domain, functionally connected standing waves of neural activity and propagating vasomotor waves over the brain cortex are the primary contributors of ISF BOLD signal variance (Bolt et al., 2022). These two mechanisms share overlapping frequencies, making them difficult to disentangle. However, the spatial ICA detects only stationary signal sources emphasizing the neuronal RSN effects more on the coupling patterns in relation to NF domains. The PAC coupling strength was higher with NF components, and it increased in sleep. An especially strong PAC was detected in the perisylvian CSF4 component, where it increased three to fourfold in sleep compared to waking (see Fig. 3b). Even though the PAC and ISF power increased in sleep, (see Fig. 2-3), the TE strength of prediction was not altered during sleep (see Fig. 4b)

Predictive TE analysis further indicated that ISF RSN activity almost exclusively drove NF component dynamics during wakefulness, where also CSF compartments predicted venous sinus changes (see Fig. 2e). This result is consistent with classical activation hyperemia, whereby cortical RSN activity increases the local cerebral blood flow, which is reflected successively into CSF and then venous sinuses sources due to volume displacement (Fultz et al., 2019; Väyrynen et al., 2024; Hauglund et al., 2025). During sleep, these relationships became more complex and were no longer coordinated as exclusively top-down cortical sources; NF sources during sleep started to show bottom-up effects on the RSN. During sleep, the causal dynamics were better characterized with chained interactions between RSN and NF compartments, implying a reorganization of the hydrodynamic driving effects relative to the waking state. The increasing ISF power in NREM 2 seems to be connected to the increase in sinus S3 and S4 towards left FPN and SN1 RSNs, (see Fig. 2e). We suppose that the increased pulsation power in veins could induce up-stream hydrodynamic effects along (peri)venous structures in the RSNs.

The data dimension reduction with ICA to 20 spatially-independent components enabled the use of surrogate statistical testing in contrast to raw coupling values in the voxel level analysis. This helped us to assess whether observed coupling patterns were genuine or arising by chance in consideration of the massive multiple comparison issue arising with an exhaustive search over the whole frequency space. Indeed, significant coupling frequency ranges assessed at the component level differed from voxel level results. For example, an apparent ISF–Card coupling observed at the voxel level may have spuriously resulted from high autocorrelation of the propagating arterial impulses, rather than representing a genuine coupling. In contrast, surrogate statistics also revealed a cardiorespiratory coupling pattern that was not evident at the voxel level. Notably, this proved to be the most stable coupling pattern across all arousal states, while ISF-related couplings were more apparent during NREM-2 sleep. PAC magnitudes depend on signal powers, and sleep increases ISF power, suggesting that the elevated PAC levels observed during sleep (see Figs. 1, 3) could also relate to an increased power level.

### Sleep-dependent amplification of vasomotor dynamics

The sleep-associated increase in ISF power across both RSN and NF components aligns with prior evidence in mice and humans that vasomotor oscillations increase during NREM sleep (van Veluw et al., 2020; Helakari et al., 2022; Bojarskaite et al., 2023; Väyrynen et al., 2024; Hauglund et al., 2025). The increase in relative ISF power with respect to total signal power (see Fig. 2c) suggests that this was not due to elevation in overall signal power levels, but rather reflects a true increase in ISF bandwidth, an interpretation also supported by the absolute power results (see Fig. 2b). Notably, some ICs exhibited very low relative ISF power levels, largely due to high cardiorespiratory pulsatility, which significantly reduced ISF power relative to total signal power levels.

In mice, decreased locus coeruleus neuron activity and associated ∼0.02-0.03 Hz fluctuations in cortical norepinephrine levels during sleep entrain vasomotor oscillations in vascular walls, which enhance perivascular CSF flow (Osorio-Forero et al., 2021; Kjaerby et al., 2022). We have previously shown that MREG ISF signal power levels globally increase during sleep (Helakari et al., 2022; Väyrynen et al., 2024). Our present findings extend this concept: declining NE levels during NREM-1 and NREM-2 sleep are permissive to increased vasomotor activity within arterial walls, which manifests as increased ISF power in arterial IC components, but also in downstream venous components, which dominate the increases in BOLD signal. This likely implies increased vascular wall oscillations arising from low frequency fluctuations in locus coeruleus noradrenergic signaling. However, the interesting upstream effects from NF towards RSN requires more in depth analysis in the future.

### Functional implications for neurofluid circulation

Beyond modulatory PAC effects, we hypothesize that transient phase alignment among vasomotor, respiratory, and cardiac pulsations could create constructive interference, and thereby amplify pressure gradients to promote CSF transport. Conversely, desynchronization induced by changes in vascular stiffness or brain activity state could result in destructive interference, which could diminish CSF bulk flow. This conceptual framework could provide a mechanistic link between sleep, pulsation synchrony, CSF flow, and glymphatic function, with implications for pathologies such as Alzheimer’s disease, which are characterized by altered brain pulsatility.

Together, these findings reveal that human brain pulsations are functionally driven across distinct slow and cardiorespiratory frequency bands forming a tightly coordinated hierarchical system in which slow hemodynamic changes coordinate faster oscillatory events from independent cortical sources towards NF compartments. The observed strengthening of this coupling during NREM sleep supports the notion of increased directional drive of CSF flow during sleep in relation to enhanced brain solute transport during sleep.

## Materials and Methods

### Study design and measurements

Our study group consisted of N=23 healthy controls (13 females, 10 males, mean e age 27.0 years), who had been investigated in our prior studies (Helakari et al., 2022, 2023). The volunteers participated in two separate fMRI scanning sessions, each lasting about one hour: one scan measured wakefulness and the other aimed to capture sleep episodes. Approximately half (N=13) of the subjects had undergone one night of sleep deprivation prior to scanning to facilitate rebound sleep. Their sleep deprivation had been verified by recordings obtained the prior night using smart rings (Oura Health Oy). Written informed consent had been obtained from all participants in accordance with the Declaration of Helsinki. Our study was screened and approved by the Regional Ethics Committee of the Northern Ostrobothnia Hospital District. In synchrony with the fMRI, we obtained EEG recordings to delineate awake epochs from sleep states and to separate NREM-1 and -2 sleep stages. Sleep stage classification was performed manually in 30-second EEG époques by experienced neurophysiologists (J.P., M.K.) according to the criteria defined by American Academy of Sleep Medicine (AASM).

We used a 3T Siemens MAGNETOM Skyra scanner with 32-channel head coil to capture the structural and functional MREG image series. The imaging parameters for structural 3D MPRAGE scans were: TR=1900 ms, TE= 2.49 ms, FA=9°, FOV=240 mm and slice thickness of 0.9 mm. An MREG-sequence was used to measure functional datasets using the following parameters: TR=100 ms, TE=36 ms, FA=5°, FOV=192 mm. The MREG sequence deliberately undersamples k-space by reaching a 10 Hz sampling frequency with voxel size of 3 mm (Assländer et al., 2013). The crusher gradient of 0.1 was optimized for physiological signals sources, which also mitigates slow drifts and stimulated echoes. The image reconstruction used L2-Tikhonov regularization with a lambda value of 0.1, as determined by the L-curve method (Hugger et al., 2011). To reduce B0-field artifacts, we used dynamic off-resonance correction in k-space.

Electroencephalography was recorded using the GES 400 EEG setup (Magstim EGI), consisting of a direct current-coupled amplifier (Net Amps 400), and 256-electrode MRI compatible electrode net (HydroCel Geodesic Sensor MR net). We used electrode ‘Cz’ located at the vertex of the head as the reference electrode. The EEG recordings were conducted with sampling rate of 1 kHz, except for three sleep and five awake scans, where 250 Hz sampling was used erroneously.

### Data preprocessing steps

MREG image processing was performed using the FSL library (Jenkinson et al., 2012) (Functional Magnetic Resonance Imaging of the Brain software library). We used a cutoff frequency of 0.008 Hz to high-pass filter the datasets, and then applied MCFLIRT motion correction and further despiking (Cox, 1996) to eliminate artifactual spikes. The structural T1 images were used to spatially normalize functional images into the MNI152 standard space. Two-minute length segments of wakefulness and NREM1-2 sleep were identified for each participants using the EEG-based sleep scores and utilized in further analysis. The NREM-1 dataset consisted of N=20 (10 females, 10 males, average age: 26.2 years), and the NREM-2 set consisted of N=14 (7 females, 7 males, average age 27.8 years).

EEG measurements were also preprocessed for MR-induced artifacts. Ballistocardiographic and gradient switching related artifacts were removed using template subtraction (Allen et al., 1998, 2000) in Brain Vision Analyzer (Brain Vision Analyzer v.2.1, Brain Products).

### Analysis of independent components

Group ICA was calculated with the FSL MELODIC package (Beckmann et al., 2009) using a temporal concatenation approach and including the full-length scans across awake and sleep recordings (See Fig 2a, d). To circumvent the tendency for components to split into subcomponents in higher model orders (Abou-Elseoud et al., 2010), we reduced the data dimensionality to 20 components using PCA. We applied the variance normalization option to ensure equal weighting across voxels.

Subsequently, a neuroradiologist (VK) visually inspected and labeled the template networks (Kiviniemi et al., 2003, 2009), whereupon we computed subject-specific networks through FSL’s dual regression method (Beckmann et al., 2009). This procedure involved fitting template networks to the subject-specific fMRI datasets, resulting in a set of subject-specific independent components corresponding to the templates.

We calculated spectral powers extending from 0.01 to 5 Hz (see Fig. 2b, c) using a wavelet convolution approach (Cohen, 2014), and then obtained instantaneous power estimates by squaring the complex magnitude results 𝑃 = |𝑧|^2^. We further estimated the ISF bandpower (0-01-0.1 Hz) using integral approximation and rectangular windowing. Instead of focusing exclusively on absolute ISF powers, we obtained relative power levels by dividing the ISF powers by total signal power to avoid effects of altered overall signal power levels.

### Estimating frequency specific coupling ranges

To quantify cross-frequency coupling across the whole frequency range, we used phase-amplitude coupling (PAC), which quantifies the relationship between the phase of one frequency and power in another (Canolty and Knight, 2010; Cohen, 2014) (See Fig. 1, 3). We calculated PAC with wavelet frequencies across the 0.01 to 5 Hz range using 300 linear increments. Wavelet convolution with complex Morlet wavelets (N=6) was used to filter the signals at each frequency step. From the convolution results, we further extracted the phase and power corresponding to each frequency bin. PAC was defined as 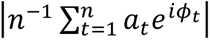, where 𝑎 is the power at time point 𝑡, 𝑖 is the imaginary operator, 𝜙 is phase angle in radians, and 𝑛 is the number of time points (Cohen, 2014). We iterated PAC calculations for each (*f_phase_,f_power_*) frequency combination, producing two-dimensional coupling estimates. A random sampling approach (N=3405) without replacement was used to assess the whole brain PAC, utilizing 5% voxels of the whole brain volume (See Fig 1c).

PAC patterns were further assessed across the frequency space with ICs (See Fig. 3). Having obtained reduced data dimensionality, we utilized surrogate data (Ns=100) to evaluate the statistical significance of each *f_phase_,f_power_* point. This approach revealed PAC clusters, of which we analyzed in more detail. For each subject, we extracted their individual cardiac and respiration frequencies and static ISF and SF bands, to study their several coupling patterns (ISF-SF, ISF-Resp and Resp-Card) in greater detail.

### Assessing directional dependencies

To calculate information transfer in the ranges with coupling, we used the phase transfer entropy (TE) approach (Lobier et al., 2014). This process involved first estimating the analysis lag, before proceeding to the actual TE estimation. Here, TE was estimated between the phase of the slow oscillations and the phase of the amplitude envelope of the fast oscillation (Vanhatalo et al., 2004; Palva et al., 2005; Väyrynen et al., 2023).

As in PAC magnitude analysis, we used static frequency bands for ISF and slow frequency bands. For the respiratory and cardiac bands, we used the individual peak frequencies as center frequencies for finite impulse response (FIR) band pass filters, using a filter order of N=10^3^ and band width of 0.2 Hz for respiratory and 0.1 Hz for cardiac frequency bands. Application of the mirroring technique with recursive filtering ensured accurate phase estimates.

We first estimated the shared information between the two pairs of variables using time-delayed mutual information (TDMI) analysis. In this approach, we assessed lags ranging from –2 to 2 seconds in 0.1 s step size. The lag value maximizing mutual information was used in the estimation of phase TE (Lobier et al., 2014), which proceeded by applying the Hilbert transform to acquire the analytical signals, using the band pass filtered signals. For the slow oscillation, phase was extracted from the analytical signal. For the fast oscillation, we first calculated the analytical envelope, which we then filtered to the corresponding slow frequency band, followed by extraction of phase (see Fig 4a).

We then proceeded to calculate TE between the pair of phase time-series with the TDMI analysis lag estimates (see Fig 4b). Here, we used a discrete estimator, essentially binning the data to represent the state space. Phase TE was then calculated within each IC and in the three identified coupling ranges in both interaction directions. Pairwise TE was then formulated into its directional form, defined as a difference *ΔTE=TE(x→y)-TE(y→x)*.

The same approach was taken to estimate the connectivity between the RSNs and NF components (see Fig 2e). Here, we utilized the ISF filtered subject-specific networks to focus specifically on the lower end of the spectrum. As before, we likewise began by estimating the analysis lag that maximized mutual information between all combinations of the ICs. To this end, we used TDMI with possible lags ranging from ±5 s, and then proceeded to calculate TE between all IC combinations using the optimal analysis lag.

### Statistical analysis

We did not make a priori calculation of sample size based on statistical methods. For all statistical tests in this exploratory study, we used an alpha level of 0.05 to infer statistical significance. The differences in relative ISF powers (Fig. 2c) were compared between the awake-NREM-1 and awake-NREM-2 contrasts using Wilcoxon rank-sum test, with the null hypothesis that the median ISF powers are equal between the arousal states. To control false positive results, we used Benjamini-Hochberg false discovery rate (FDR) correction (Benjamini and Hochberg, 1995). A similar approach was taken to test the raw PAC values of IC components (Fig. 3b). We used the sign test and FDR-correction to assess if the information transfer has a net directionality in the three observed coupling ranges (Fig. 4b).

Moreover, we utilized surrogate statistics (Theiler et al., 1992; Lachaux et al., 1999; Lobier et al., 2014) to perform model free, empirical statistical analysis (Fig 2e,3a, 4). The null hypothesis distributions were built using Ns=100 surrogates, where each surrogate signal was constructed using the time-shift method. In this approach, the original signal is split and reattached from a random time-point, which preserves the signal autocorrelation structure of the original data, but removes any linear correlations.

## Acknowledgments

This study was funded by Research counsil of Finland: TERVA grants 1-2, 275342, 338599, 314497, 335720 (V.Ki), 360508 (J.K), The EU Joint Programme – Neurodegenerative Disease Research 2022-120 (V.Ki), Jane & Aatos Erkko Foundation: 1, 210043 (V.Ki.), VTR grants from Oulu University Hospital (V.Ki, V.Ko, J.K). Instrumentarium Science Foundation (T.V, J.T, M.J),, The Finnish Medical Foundation (V.Ki, M.J), Finnish Brain Foundation (V.Ki, L.R, J.T), Pohjois-Suomen Terveydenhuollon tukisäätiö (H.H, V.Ko), Tauno Tonning Foundation (L.R), Juhani Aho Foundation for Medical Research (L.R), Uniogs/MRC Oulu DP-grant (H.H), Emil Aaltosen Säätiö (H.H, L.R, M.J). Finnish Cultural Foundation (M.J), Orion Science Foundation (M.J), The Paolo Foundation (M.J). We thank Adj. Prof. Paul Cumming of Queensland University of Technology for feedback and comments on the manuscript. We further thank CSC – IT Center for Science Ltd., Finland for providing high-performance computational resources used in this study. We thank Oura Health and Hannu Kinnunen for their collaboration.

## Supplementary Information

**Supplementary Table 1.** List of abbreviations used in naming of independent components.

|  |  |
| --- | --- |
| RSN | Resting state networks |
| SN1 | Salience network |
| SN2 | Salience network |
| DMNa | Default mode network, anterior |
| DMNp | Default mode network, posterior |
| FPNI | Fronto-parietal network, left |
| FPNr | Fronto-parietal network, right |
| SMN | Sensorimotor network |
| DAN | Dorsal attention network |
| Visl | Lateral visual network |
| Vism | Medial visual network |
| NF | Neurofluid components |
| S1 | Sinus sagittal, anterior |
| S2 | Sinus sagittal, posterior |
| S3 | Sinus sagittal |
| S4 | Sinus sagittal |
| S5 | Venous component |
| CSF1 | Cerebrospinal fluid component |
| CSF2 | Cerebrospinal fluid component |
| CSF3 | Cerebrospinal fluid component / arterial component |
| CSF4 | Cerebrospinal fluid component / arterial component |
| Art | Arterial component |

**Supplementary Table 2.** Relative infra-slow (ISF) power of independent components. Median ISF powers relative to total signal power levels are presented along with corresponding interquartile range (IQR) during wakefulness (A) and sleep states (N1/N2). The statistical outcomes of Wilcoxon rank-sum tests across arousal states (A-N1, A-N2) are shown for each component, including the *Z*-statistics and corresponding FDR-adjusted *p*-values.

|  |  | SN1 | SN2 | DMN_a | DMN_p | FPN_l | FPN_r | SMN | DAN | Vis_l | Vis_m |
| --- | --- | --- | --- | --- | --- | --- | --- | --- | --- | --- | --- |
| A | *P (%)* | 13.17 | 7.29 | 4.68 | 15.05 | 18.03 | 20.20 | 14.06 | 19.91 | 6.85 | 12.20 |
|  | *IQR (%)* | 10.70 | 9.27 | 6.61 | 9.32 | 13.50 | 9.21 | 16.26 | 14.75 | 9.95 | 20.61 |
| N1 | *P (%)* | 13.64 | 13.34 | 5.13 | 19.26 | 19.58 | 13.51 | 18.93 | 17.43 | 12.56 | 33.74 |
|  | *IQR (%)* | 13.78 | 11.82 | 2.88 | 16.32 | 11.71 | 9.42 | 11.88 | 19.16 | 9.20 | 24.50 |
| N2 | *P (%)* | 15.12 | 23.54 | 3.87 | 30.46 | 19.41 | 15.88 | 24.09 | 28.73 | 15.03 | 48.44 |
|  | *IQR (%)* | 22.27 | 12.83 | 2.00 | 15.43 | 6.95 | 11.25 | 20.47 | 15.78 | 12.07 | 20.97 |
| A-N1 | *Z* | 0.305 | -2.068 | 0.141 | -0.940 | 0.752 | 2.444 | -0.987 | 0.540 | -2.115 | -2.021 |
|  | *p.adj* | 0.800 | 0.077 | 0.888 | 0.496 | 0.549 | 0.060 | 0.496 | 0.654 | 0.077 | 0.079 |
| A-N2 | *Z* | -0.611 | -3.367 | 1.143 | -3.586 | -0.172 | 0.893 | -2.020 | -1.394 | -2.991 | -4.181 |
|  | *p.adj* | 0.602 | 0.004 | 0.337 | 0.003 | 0.909 | 0.438 | 0.067 | 0.233 | 0.008 | 0.001 |

|  |  | S1 | S2 | S3 | S4 | S5 | CSF1 | CSF2 | CSF3 | CSF4 | Art |
| --- | --- | --- | --- | --- | --- | --- | --- | --- | --- | --- | --- |
| A | *P (%)* | 8.04 | 2.77 | 1.18 | 10.40 | 12.25 | 3.23 | 2.18 | 4.81 | 1.66 | 0.43 |
|  | *IQR (%)* | 8.06 | 4.92 | 2.62 | 11.38 | 15.11 | 3.41 | 1.88 | 5.04 | 2.56 | 0.64 |
| N1 | *P (%)* | 7.81 | 8.98 | 2.12 | 23.25 | 19.27 | 2.75 | 5.03 | 8.76 | 3.98 | 1.13 |
|  | *IQR (%)* | 9.10 | 12.26 | 3.87 | 17.48 | 22.31 | 3.25 | 5.62 | 7.59 | 3.15 | 1.21 |
| N2 | *P (%)* | 12.35 | 14.54 | 3.97 | 24.16 | 29.90 | 3.45 | 6.96 | 10.05 | 4.58 | 1.01 |
|  | *IQR (%)* | 11.21 | 14.34 | 5.19 | 33.16 | 26.00 | 2.44 | 5.33 | 8.55 | 2.09 | 1.13 |
| A-N1 | *Z* | -0.752 | -2.397 | -2.256 | -2.256 | -1.715 | 0.728 | -3.760 | -2.632 | -2.303 | -3.736 |
|  | *p.adj* | 0.549 | 0.060 | 0.060 | 0.060 | 0.144 | 0.549 | 0.002 | 0.057 | 0.060 | 0.002 |
| A-N2 | *Z* | -0.986 | -2.865 | -2.208 | -2.208 | -3.085 | -0.016 | -3.492 | -2.865 | -2.302 | -3.210 |
|  | *p.adj* | 0.405 | 0.009 | 0.045 | 0.045 | 0.007 | 0.988 | 0.003 | 0.009 | 0.043 | 0.005 |

**Supplementary Table 3.** Phase-amplitude coupling (PAC) differences across resting state networks (RSN) and neurofluid (NF) components. Differences between RSNs and NF were assessed using the signed rank test for the three coupling patterns and across arousal states. *Z*-values and false discovery rate (FDR) adjusted *p*-values are shown.

|  |  |  |  |  |
| --- | --- | --- | --- | --- |
| **ISF-SF** |  |  |  |  |
| Awake | 0.096 | 0.233 | -4.106 | 1.21E-04 |
| NREM-1 | 0.194 | 0.278 | -3.883 | 1.55E-04 |
| NREM-2 | 0.231 | 0.480 | -3.296 | 9.82E-04 |
| **ISF-Resp.** |  |  |  |  |
| Awake | 0.070 | 0.173 | -4.106 | 1.21E-04 |
| NREM-1 | 0.129 | 0.218 | -3.883 | 1.55E-04 |
| NREM-2 | 0.155 | 0.335 | -3.296 | 9.82E-04 |
| **Resp.-Card.** |  |  |  |  |
| Awake | 0.021 | 0.086 | -4.197 | 1.21E-04 |
| NREM-1 | 0.024 | 0.109 | -3.920 | 1.55E-04 |
| NREM-2 | 0.026 | 0.126 | -3.296 | 9.82E-04 |

**Supplementary Table 4.** Phase-amplitude coupling (PAC) differences across arousal states. PAC median and interquartile ranges associated with each coupling pattern and arousal state. We tested pairwise differences between states (A-N1, A-N2, N1-N2) using the two-sample Wilcoxon rank sum test. Associated *Z*-values and false discovery rate (FDR) adjusted *p*-values are shown.

|  | *N* | *Median* | *IQR* | *Test* | *Z* | *p<sub>adj</sub>* |
| --- | --- | --- | --- | --- | --- | --- |
| **ISF-SF** |  |  |  |  |  |  |
| Awake | 23 | 0.174 | 0.172 | A-N1 | -1.546 | 0.215 |
| NREM-1 | 20 | 0.230 | 0.309 | A-N2 | -2.834 | 0.021 |
| NREM-2 | 14 | 0.356 | 0.277 | N1-N2 | -1.382 | 0.215 |
| **ISF-Resp.** |  |  |  |  |  |  |
| Awake | 23 | 0.120 | 0.117 | A-N1 | -1.619 | 0.215 |
| NREM-1 | 20 | 0.172 | 0.265 | A-N2 | -2.834 | 0.021 |
| NREM-2 | 14 | 0.261 | 0.221 | N1-N2 | -1.207 | 0.256 |
| **Resp.-Card.** |  |  |  |  |  |  |
| Awake | 23 | 0.059 | 0.048 | A-N1 | -0.451 | 0.652 |
| NREM-1 | 20 | 0.063 | 0.044 | A-N2 | -1.801 | 0.215 |
| NREM-2 | 14 | 0.075 | 0.032 | N1-N2 | -1.452 | 0.215 |

**Supplementary Table 5.** Net directionalities of the coupling patterns as measured by phase transfer entropy (TE). Median *ΔTE* values (in bits) along with interquartile ranges are shown for each coupling pattern and arousal state. A one sample sign test was used to test the directionality of suprathresholded (p<0.05) TE values. Associated sign and false discovery rate (FDR) adjusted *p*-values are shown.

|  | *Count* | *Median* | *IQR* | *Sign* | *Padj* |
| --- | --- | --- | --- | --- | --- |
| **ISF-SF** |  |  |  |  |  |
| Awake | 25 | 0.084 | 0.115 | 22 | 4E-04 |
| NREM-1 | 23 | 0.078 | 0.109 | 22 | 1E-04 |
| NREM-2 | 16 | 0.070 | 0.121 | 16 | 1E-04 |
| **ISF-Resp.** |  |  |  |  |  |
| Awake | 19 | 0.067 | 0.107 | 16 | 0.007 |
| NREM-1 | 16 | 0.101 | 0.123 | 16 | 1E-04 |
| NREM-2 | 20 | 0.076 | 0.163 | 18 | 0.001 |
| **Resp.-Card.** |  |  |  |  |  |
| Awake | 19 | 0.000 | 0.091 | 9 | 1.000 |
| NREM-1 | 13 | -0.012 | 0.176 | 6 | 1.000 |
| NREM-2 | 18 | -0.072 | 0.144 | 3 | 0.010 |

**Figure S1.**
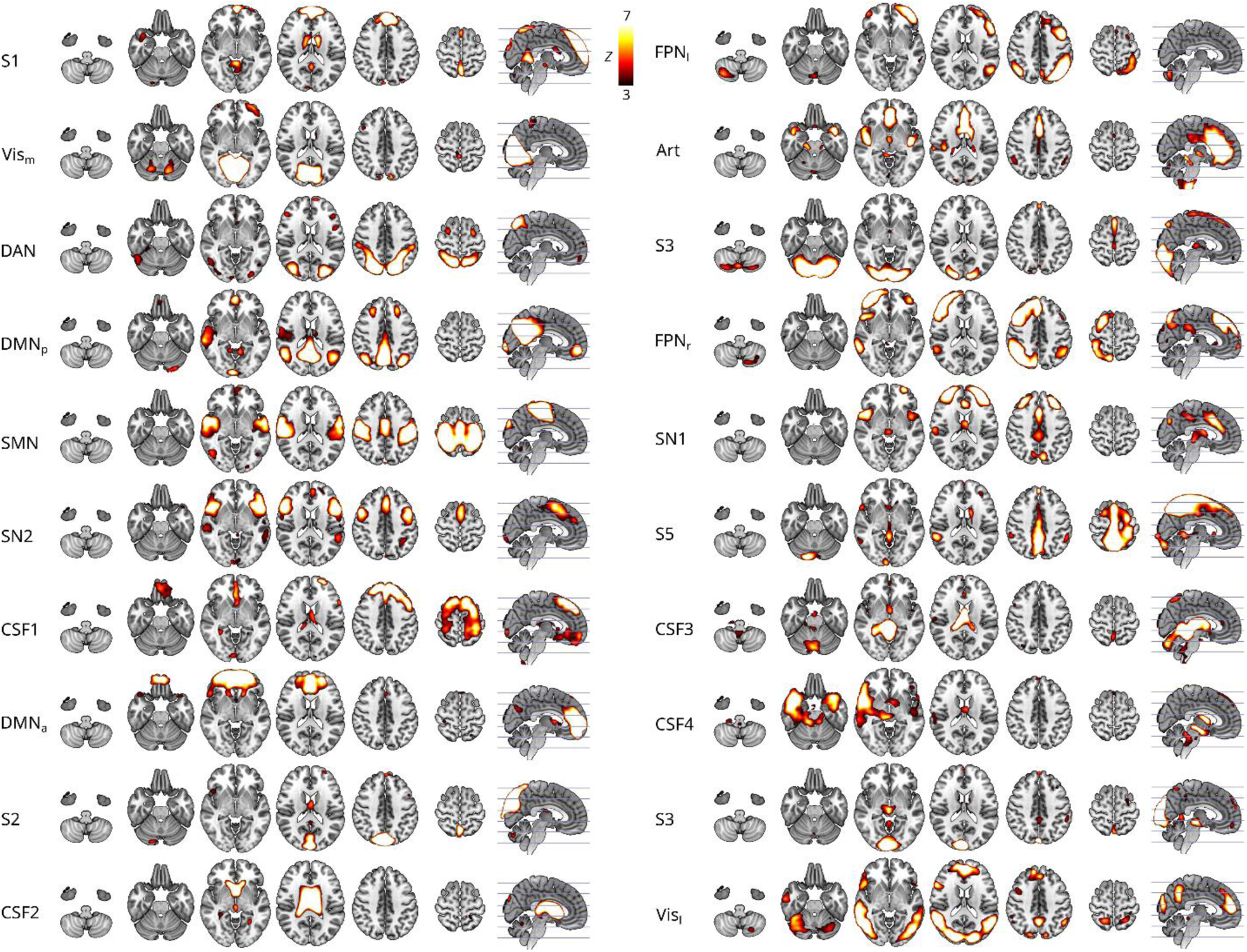
Spatial coverage (Z-values) of template networks obtained from group ICA. The networks are presented in the original order, with highest variance explained first from the left panel (top to bottom), and followed by the right panel. .

**Figure S2.**
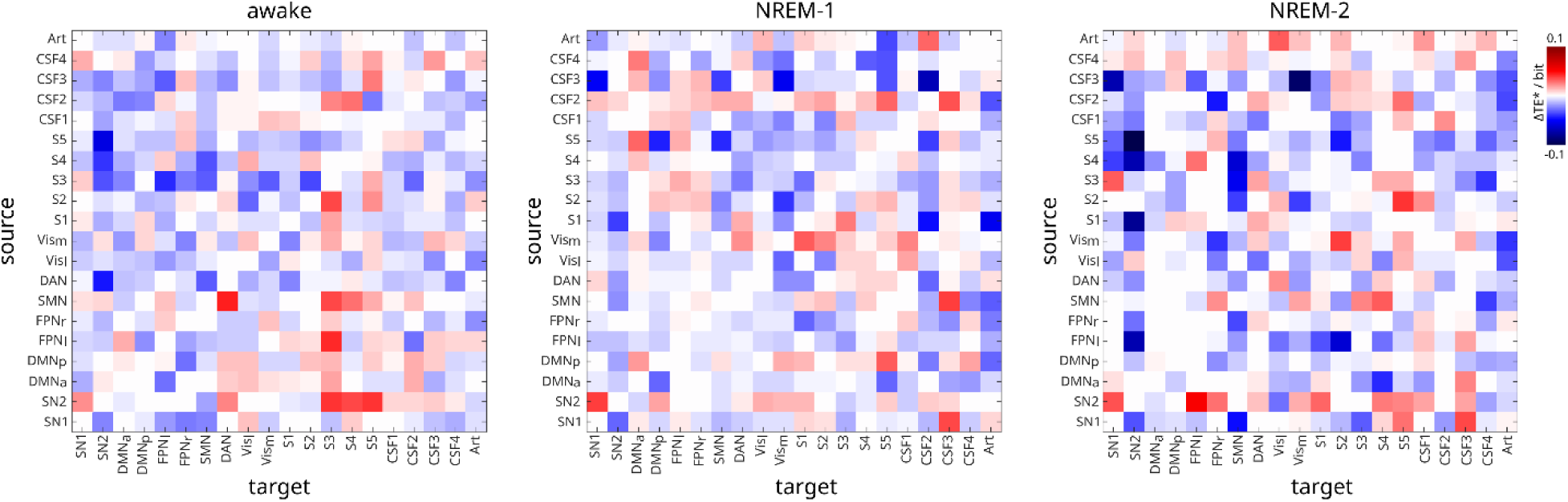
Adjacency matrix picturing directed connections between networks based on transfer entropy (TE). Surrogate thresholded (p<0.05) *Δ*TE values were averaged over all subjects to construct adjacency matrices during wakefulness and NREM-1 and -2 sleep states.

